# Small-Molecule Structure Determination and Anisotropic Displacement Analysis at Turkish Light Source

**DOI:** 10.64898/2026.04.19.719451

**Authors:** Esra Ayan, Arif Mermer

## Abstract

Single-crystal X-ray diffraction remains one of the most direct and reliable techniques for clarifying the three-dimensional structures of small molecules; however, its wider use in developing research settings has historically been limited by access to advanced instrumentation. Here, we consider the performance of the in-house diffractometer, Turkish Light Source, for small-molecule structure determination using three rhodanine-derivative compounds. Diffraction data were collected, processed, and followed by full-matrix least-squares refinement as a user-friendly pipeline. The compounds were successfully resolved in the triclinic space group P–1 and refined to chemically reasonable models, although notable differences in data quality and refinement parameters were observed. Compounds 1 and 2 produced the most robust and internally coherent structure, whereas compound 3 displayed refinement tribulations. These might be attributed to the intrinsic structural disorder of c-5b, analogous to polymorphic perversity in higher Z′ phase, likely due to the presence of dissymmetric molecules within the asymmetric unit (Z′ = 2), rather than empirical limitations. Anisotropic displacement parameters were systematically computed by atom-resolved Ueq factors and anisotropy index. The combined analyses reveal that structural ambiguity of c-5b is largely governed by localized maxima in atomic displacement (up to 0.29 Å^2^ in Ueq with 6.67 anisotropy) rather than by global disorder, caused by the fluorinated aryl moiety of c-5b. These findings indicate that the in-house SCXRD system, when coupled with our user-friendly downstream pipeline, can yield reliable structural data for small molecules.

Brief video tutorials and detailed SOPs have been provided in the [Tutorials](https://drive.google.com/drive/u/0/folders/1AZDcdPPFMRBdK8JDj-qT-VxyR34HD1Dp) folder, including [CrysAlisPro](https://drive.google.com/drive/u/0/folders/1s-44-xNMvVWSmhFIAUK-Rccy_iiBmpuP) and [Olex2](https://drive.google.com/drive/u/0/folders/12kYXxO6c6LXJuTb-Iblfiiq7gClT3UdK) tutorials, as well as are easily accessible for users.

## 1. Introduction

The synthesis of small organic molecules, particularly heterocyclic compounds with potential pharmacological relevance such as rhodanine derivatives, remains an important area of modern medicinal chemistry [1]. Compounds of this type have been related to a wide range of biological activities, including anticancer, antimicrobial, antioxidant, and enzyme-inhibitory effects, and therefore continue to draw interest in structure-based chemical research [2]. Therefore, accurate structural determination is essential for understanding molecular conformation, intermolecular interactions, and structural characteristics that may influence biological properties. Although spectroscopic methods provide valuable information regarding composition and functional-group identity, single-crystal X-ray diffraction (SCXRD) remains the most direct method for determining the three-dimensional structure of small molecules in the crystalline state at atomic resolution [3].

An ongoing practical limitation in SCXRD has been the need for suitable single crystals and for instrumentation capable of collecting high-quality diffraction data from small or weakly diffracting crystalline [3]. However, recently, advances in X-ray sources, optics, goniometer design, and detector technology have improved the performance of laboratory-based diffractometers [4], [5]. Namely, home-source instruments can now provide data quality acceptable for standard high-resolution structure determination in numerous small-molecule applications. In Türkiye, the installation of the Rigaku XtaLAB Synergy system (“Turkish Light Source”) in 2022 marked an important step in developing a local crystallographic facility by enabling high-quality diffraction measurements without dependence on external large-scale facilities [4].

Here, the present study aimed to collect, for the first time, SCXRD data for three synthesized small organic molecules, compound-5a, compound-5p, and compound-5b, and to determine their three-dimensional atomic structures using the newly installed Turkish Light Source (referred to as TLS) system. We present a pipeline here, ***(i)*** diffraction data were processed in CrysAlisPro, and ***(ii)*** structure determination and refinement were performed in Olex2. Apart from standard crystallographic refinement, ***(iii)*** anisotropic displacement tensors were extracted from the refined *.cif or *.pdb files and the parameters were evaluated using Ueq = (U11 + U22 + U33) / 3 together with an anisotropy index defined as max(U11, U22, U33) / min(U11, U22, U33), enabling comparative evaluation of the anisotropic displacement parameters (referred to as ADPs) and atom-resolved displacement factors (referred to as Ueq) of the three structures [6]. This combined technique was used to examine how differences in substituents may relate to crystal packing disorder, refinement ambiguity, and localized displacement maxima within the final structures.

## 2. Materials and methods

### 2.1. Materials

Compounds 5a, 5p, and 5b were synthesized via a one-pot, four-component reaction involving the condensation of the corresponding amine, carbon disulfide, ethyl bromoacetate, and an aromatic aldehyde under basic conditions (triethylamine) in aqueous medium. The reaction proceeds through the in-situ formation of an N-substituted rhodanine (thioxothiazolidinone) intermediate, followed by a base-catalyzed Knoevenagel condensation with the aldehyde to afford the target (Z)-5-arylidenerhodanine derivatives. While compounds 5a and 5p were obtained using 4-(3-aminopropyl)morpholine as the amine component, compound 5b was synthesized analogously using 3,5-bis(trifluoromethyl)benzylamine and 3,5-dimethylbenzaldehyde under ultrasonic conditions. The protocol can be performed under conventional, microwave, or ultrasonic irradiation, providing the desired products with good yields with minimal work-up [2], [7], [8].

### 2.2. Method

#### 2.2.1 Instrumentation

SCXRD data collection was performed at the Turkish Light Source using a Rigaku XtaLAB Synergy diffractometer implemented with a HyPix-Arc 150° hybrid photon-counting detector [4]. Crystals were mounted on MiTeGen loops and centered on a kappa goniometer under a collimated Cu Kα X-ray beam generated by a microfocus rotating anode source. Data collection was performed using the CAP suite, which enables automated crystal centering, strategy calculation, and data collection under standard single-crystal operational mode (Oxford Diffraction Ltd).

#### 2.2.2 Sample delivery and single crystal X-ray data collection

Crystallines were prepared on glass slides and looked under a compound light microscope to identify well-formed crystals suitable for diffraction. Selected crystals were manually harvested using MiTeGen mounting loops of appropriate size and transferred onto magnetic sample pins. The mounted crystals were subsequently fixed on the goniometer head and centered in the X-ray beam using the optical alignment system. SCXRD data were collected using a Rigaku XtaLAB Synergy diffractometer implemented with a HyPix-Arc 150° hybrid photon-counting detector [4]. The crystal was positioned at the intersection of the incident X-ray beam and the detector axis, where diffraction was recorded under a monochromatic Cu Kα radiation source generated by a microfocus rotating anode. Data collection was performed using the CAP suite, which enables automated crystal centering, screening, strategy calculation, and frame collection under standard single-crystal conditions. Initial screening images were collected to assess diffraction quality, followed by optimized data collection strategies to ensure sufficient completeness and redundancy. Data collection parameters, including detector distance, scan width, exposure time, and total acquisition time, were adjusted for each compound (Table 1). The instrument was operated at 40 kV and 12.5 mA, with shutterless data acquisition enabled by the HyPix detector.

**Table 1.** Data collection parameters for three small molecules.

| Parameters | Compound 5a | Compound 5p | Compound 5b |
| --- | --- | --- | --- |
| Scan width |  | 0.50 |  |
| Distance | 49.00 | 39.00 | 39.0 |
| Pixel max/max cps | 25354/949975 | 48469/148301 | 34254/17110 |
| Exposure time min/max | 0.5/0.5 | 0.5/0.5 | 2.0/2.0 |
| Dose time | 3m 52s | 37m 4s | 2h 32m 32s |
| Total time | 16m 46s | 46m 18s | 2h 42m 5s |
| HyPix readouts | 13920 | 88960 | 4576 |
| kV |  | 40.4 |  |
| mA |  | 12.5 |  |
| Shutterless operation | 70.0 Hz (16bit-ZDT) | 40.0 Hz (16bit-ZDT) | 70.0 Hz (16bit-ZDT) |
| Frames/runs | 4640/48 | 4448/47 | 4576/50 |

#### 2.2.3 Structure determination and refinement

Following data collection, diffraction datasets were reduced and processed using the CAP software suite (Oxford Diffraction Ltd). The processed reflection files were used for structure solution and refinement in the Olex2 crystallographic package [9]. For three compounds, structure determination was performed using SHELXT (Intrinsic Phasing), followed by full-matrix least-squares refinement against F² using SHELXL within the Olex2 interface. Non-hydrogen atoms were refined anisotropically, while hydrogen atoms were placed in calculated positions and refined using a riding model.

***(i)*** for compound c-5a (C17H18ClFN2O2S2), hydrogen atoms were refined by fixed Uiso values set at 1.2×Ueq of their parent atoms (all C–H and CH2 groups). Secondary CH2 groups (C11–C17) were refined using riding coordinates, and aromatic/amide hydrogens (C1, C2, C6, C7) were also treated with a riding model.
***(ii)*** for compound c-5p (C17H19N2O2S4), hydrogen atoms were refined by fixed Uiso values again (1.2×Ueq for C–H, CH2, and S–H groups). Rigid-bond (RIGU/DELU-type) restraints were applied to C8, C10 atoms, and Uiso/Uaniso similarity restraints were implemented for displacement parameters of atoms within 2.0 Å. Secondary CH2 groups (C11–C17) and aromatic/amide hydrogens were refined using riding coordinates. Additionally, idealized S–H groups were refined as rotating groups.
***(iii)*** for compound c-5b (C42H30F12N2O2S4), hydrogen atoms were refined with Uiso values fixed at 1.2×Ueq for CH/CH2 groups and 1.5×Ueq for methyl groups. Extensive rigid-bond restraints were applied to fluorinated atoms (F1–F12) and selected C–F/N–C pairs. Uiso/Uaniso similarity restraints were applied to all non-hydrogen atoms within 2 Å. Secondary CH2 groups were refined with riding coordinates, while aromatic hydrogens were treated using a riding model. Methyl groups were placed in calculated positions and refined using a riding model.

All three structures were successfully indexed in the triclinic centrosymmetric space group P–1 and refined to chemically reasonable models, with refinement statistics summarized in **Table-2**. Differences in refinement quality among the datasets were observed, reflecting variations in crystal quality and structural complexity. Detailed workflows for data processing and reduction are provided in the SOP/video tutorial “data_process_CAP”, while step-by-step procedures for structure determination and refinement in Olex2 are described in “data_refinement_OLEX2”.

#### 2.2.4 Anisotropic displacement analysis

Anisotropic displacement parameters (ADPs) were extracted from the refined structural models obtained in Olex2 using the final *.cif files [9]. The tensor components (U11, U22, U33, U12, U13, U23) were parsed and analyzed to evaluate atom-resolved displacement factors across the three structures. An approximate equivalent anisotropic displacement parameter (𝑈𝑒𝑞) was calculated as 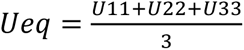, and an anisotropy index was defined as the ratio of the maximum to the minimum principal diagonal term; 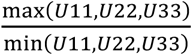 [6]. The extracted parameters were further processed using gemmi, pandas, and matplotlib libraries to generate atom-resolved displacement profiles and heatmap representations of the 𝑈𝑖𝑗 tensors. This combined analysis enabled comparative assessment of local versus global displacement maxima and facilitated identification of atoms exhibiting enhanced anisotropy or localized structural ambiguity.

## 3. Results

The compounds under the question here compounds (c-5a and c-5p) were previously reported as potent carbonic anhydrase II inhibitors obtained via a one-pot multicomponent synthesis [7], whereas c-5b has been described within a related rhodanine-based framework with therapeutic relevance [8]. Despite sharing a common rhodanine core, these compounds differ in substituent pattern and structural complexity, providing a suitable basis for comparative crystallographic analysis.

### 3.1. Small-molecule X-ray data collection, processing and data quality assessment

For data collection, a small amount of grease was first smeared to a glass slide, after which the relevant compound was placed onto the surface (**Fig. 1a**). The crystals were examined under a microscope, and a single crystal with suitable morphology was carefully selected, delivered using a MiTeGen pin, and mounted on the IGH goniometer [4]. After centering the crystal in the X-ray beam (**Fig. 1b**), diffraction data were collected (**Fig. 1c**) under the conditions summarized in **Table 1**.

**Figure 1.**
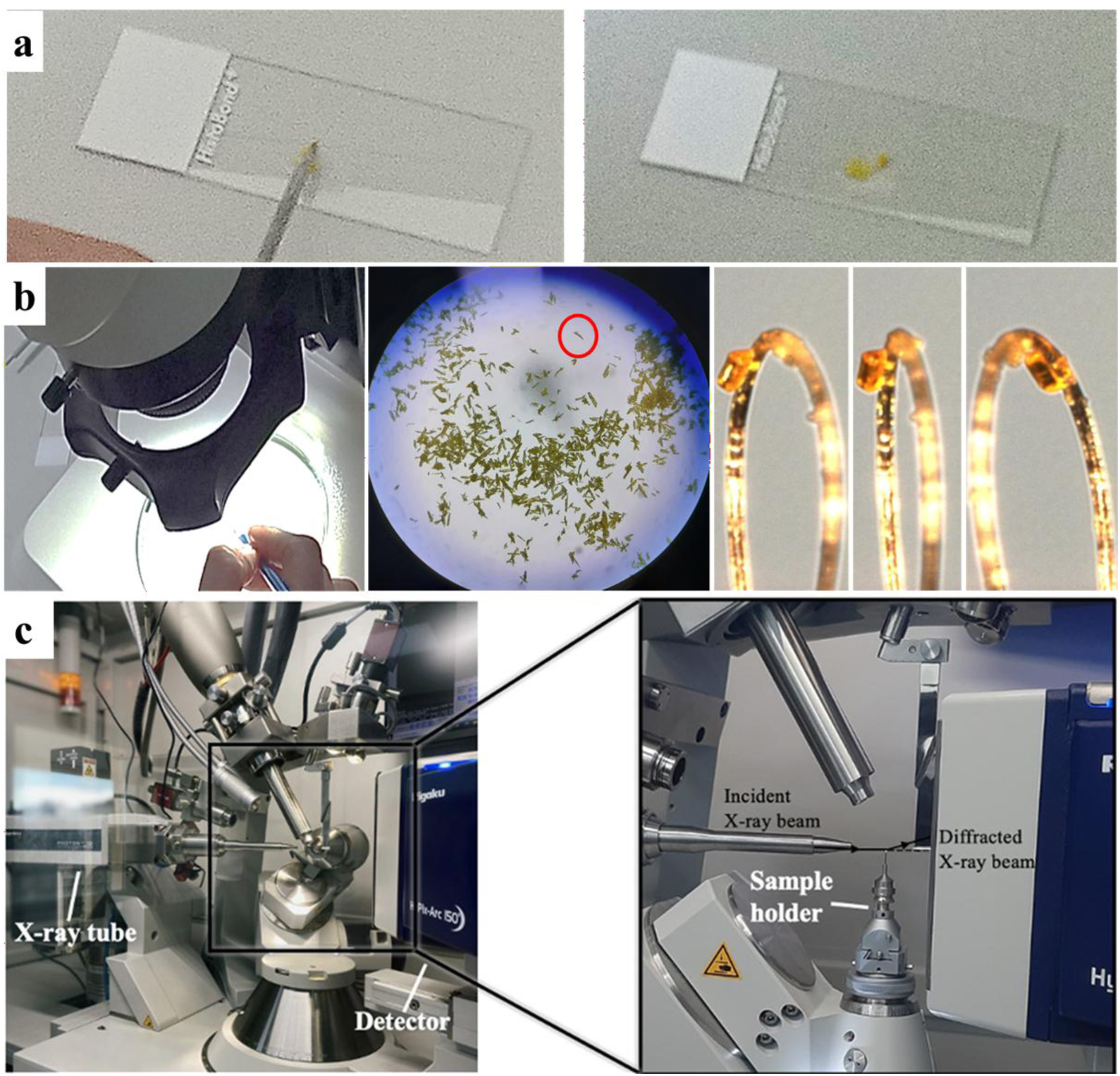
Small molecule sample delivery for diffractometer. (a) Sample preparation, (b) crystal selection and centering, and (c) mounting workflow for SCXRD data collection (panel c is adapted from Atalay et al. [4]).

Following data collection and before refinement, the diffraction datasets for those compounds were reduced by CrysAlis^Pro^ (referred to as CAP) suite. Overall, the processing statistics indicate that all three datasets were well-suited for structure determination and refinement, although their internal quality metrics reveal noticeable differs in data quality. Importantly, all three datasets were consistent with the triclinic centrosymmetric space group P-1. Among the three compounds, compound 5a (c-5a) yielded the highest-quality dataset. Processing of 4640 frames retained 12198 reflections out of 17009 tested reflections, with a low average mosaicity of 0.48. Integration and fitting were well-behaved, producing 17641 fitted reflections with no overfit or bad reflections and only 123 rejected outliers. The dataset reached 96.8% completeness at 0.80 Å and showed the most favorable merging statistics, including a low Rint value of 0.034 and a high F2/sig(F2) value of 37.3.

The compound 5p (c-5p) dataset also showed strong overall performance and was highly suited for further refinement. All 4448 collected frames were retained for processing, and three-dimensional profile analysis preserved 7384 reflections out of 19587 tested reflections, with an average mosaicity of 0.47. Integration yielded 20376 fitted reflections with no overfit or bad reflections, while 290 outliers were rejected. This dataset extended to an information limit of 0.80 Å with 92.0% completeness and gave a low Rint value of 0.048. Although slightly less internally consistent than c-5a, these statistics indicate a complete, well-resolved, and reliable dataset suitable for confident structure solution and refinement.

By comparison, compound 5b (c-5b) produced a somewhat less robust, but still interpretable, dataset. Processing of 4576 frames retained 8368 reflections out of 42706 tested reflections, with an average mosaicity of 0.50. Integration and profile fitting remained stable, yielding 42508 fitted reflections with no overfit or bad reflections, although the number of rejected outliers was higher (513). The dataset was indexed in P-1, reached 95.8% completeness at 1.00 Å. However, its relatively higher Rint value of 0.059 and lower overall F2/sig(F2) value of 13.4 suggest reduced internal consistency relative to the other two. Still, the dataset remained of enough quality to support structure determination and further refinement. Taken together, these results show that all three compounds yielded processable single-crystal diffraction data, with c-5a providing the most robust dataset, c-5p also showing strong crystallographic performance, and c-5b delivering somewhat weaker but still usable data.

### 3.2. Crystal indexing to structure refinement

After crystal centering and before data reduction, 61 images were collected for each compound to document crystal morphology and to support face indexing. Scanning of the crystals at φ = -180° and φ = 90° showed that the same compound could be followed reliably through different orientations, allowing multiple exposed faces (corresponding to hkl indices) to be identified in three dimensions (**Fig. 2ai–iii**). The diffraction images of those compounds recorded from orientations showed detectable Bragg reflections from each crystal, confirming that all three orientations were diffraction-active (**Fig. 2bi–iii**). But the patterns suggested subtle and meaningful differences in data quality; c-5a showed more clearly defined reflections, whereas c-5p and especially c-5b appeared somewhat less well resolved. The diffraction disorders were clear at the reciprocal space, but they were later supported more by the further processing (video tutorial entitled ‘data_process_CAP’) and refinement results (**Fig. 2ci–iii**).

This is apparent during refinement in Olex2 (video tutorial entitled ‘data_refinement_OLEX2’), where c-5a gave the most satisfactory structural model, followed by c-5p, while c-5b proved to be the least robust of the three. c-5a refined to R1 = 3.51%, wR2 = 9.70%, GoF = 1.089, and Rint = 3.19%, consistent with a well-behaved and internally coherent model; c-5p gave intermediate statistics, with R1 = 8.11%, wR2 = 28.77%, GoF = 1.077, and Rint = 4.82%. By contrast, c-5b showed the weakest refinement metrics, with R1 = 10.86%, wR2 = 34.41%, GoF = 1.312, and Rint = 6.14%. Taken together, these values support the ranking c-5a > c-5p > c-5b and identify c-5b as the most problematic during refinement (**Fig. 2ci–iii**).

A similar pattern was supported by the powder-diffraction comparisons generated from the Olex2 structure factors and the preprocessed data from CAP; c-5a and c-5p showed better agreement between calculated and experimental profiles than c-5b, and this was reflected in the SSE (sum of squared errors) values [10]. The SSE remained low for c-5a (**Fig. 2d**; 2.9505 × 10^-4^) and was particularly low for c-5p (**Fig. 2e**; 1.0343 × 10^-5^), whereas c-5b showed a noticeably higher value (**Fig. 2f**; 2.9564 × 10^-3^).

**Figure 2.**
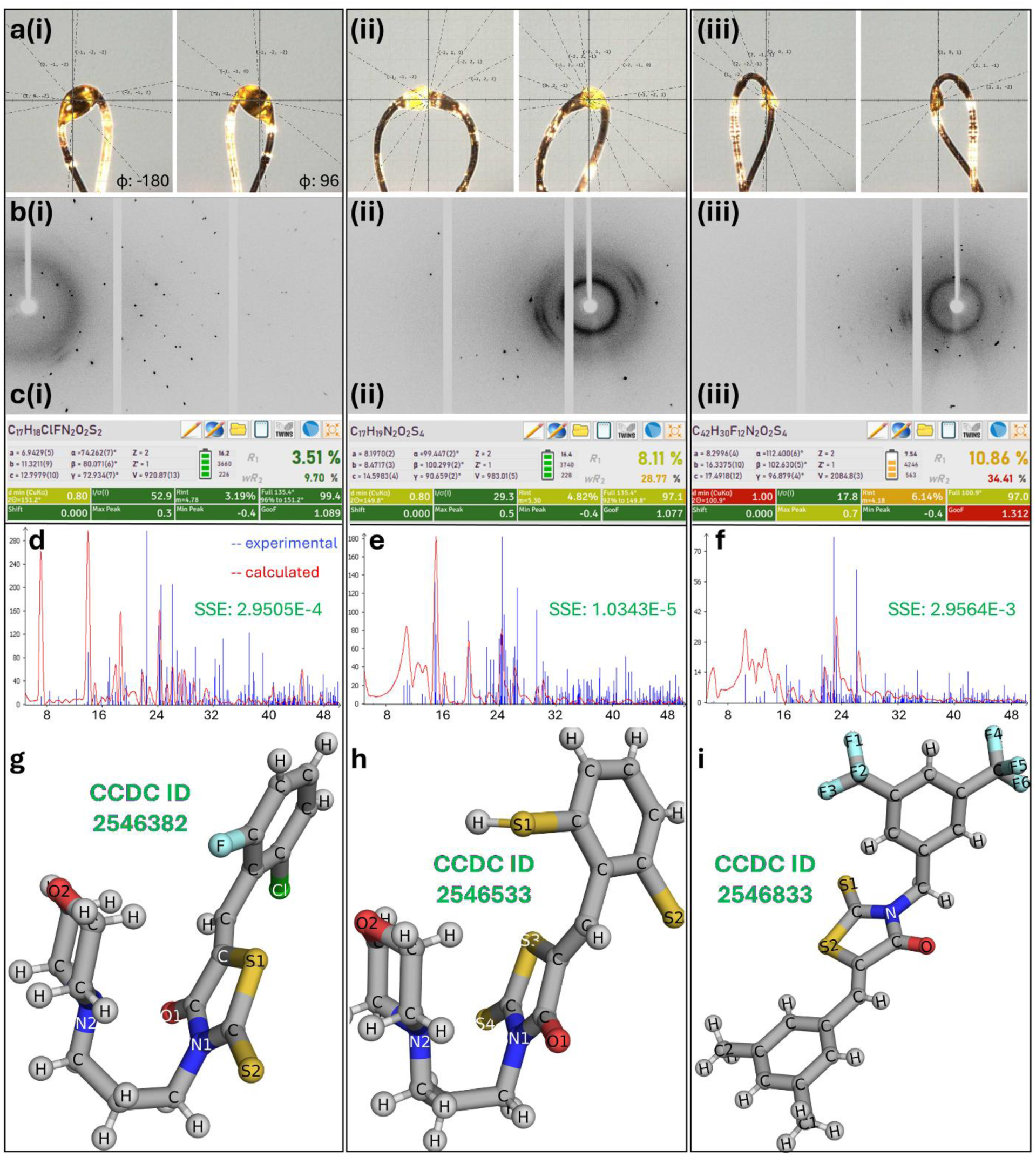
Small molecule data reduction, further refinement and comparative analysis of the datasets for three-molecules in stick mode. (a, inset i-iii) face-indexed crystal orientations at different φ angles, (b, inset i-iii) their corresponding diffraction patterns, (c, i-iii) refinement statistics obtained from Olex2 suite, (d-f) powder-pattern agreement between experimental and calculated data, and (g-i) the final refined molecular structures with associated CCDC deposition numbers. Images are generated by PyMOL.

The final refined 3D structures (*.cif) were also highlighted with clear chemical differences among the three compounds. Structure c-5a is characterized by the presence of the novel F and Cl substituents (**Fig. 2g**). Structure c-5p contains additional sulfur atoms, including the additional S sites visible in the refined model (**Fig. 2h**). Structure c-5b carries six F atoms together with two methyl groups, giving rise to the most highly substituted among the three (**Fig. 2i**). The more challenging behavior of c-5b may partly reflect its greater crystallographic complexity, since its asymmetric unit (ASU) contains two molecules of the same compound rather than one, as in the counterparts (**Fig. 2i**; here only one is shown for clarity). In sum, all three compounds yielded interpretable structures, but c-5a was the strongest, c-5p remained reliable, and c-5b was the most demanding to model and refine. Further data statistics of three compounds have been shown in **Table 2**.

**Table 2.**
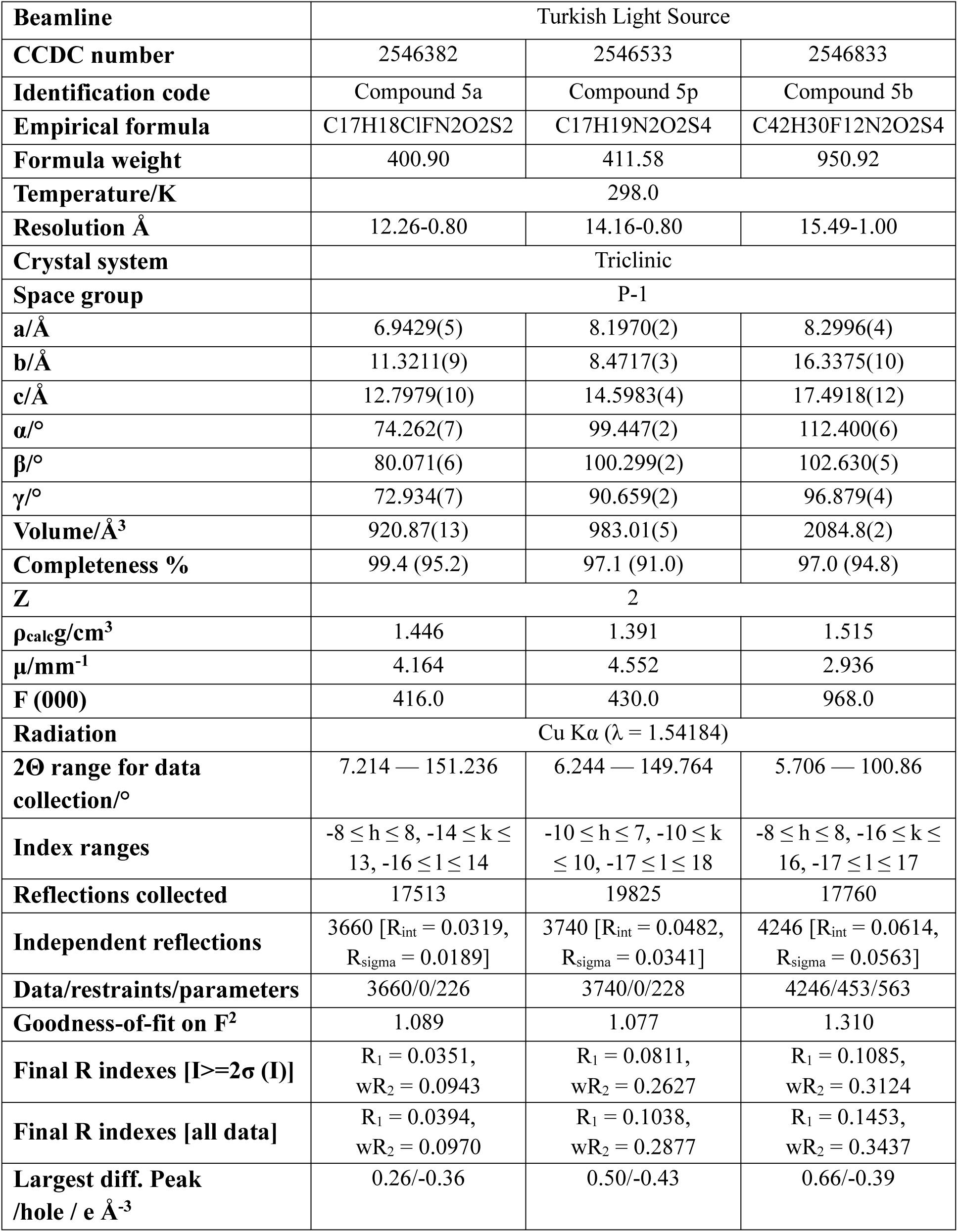
Crystal data and structure refinement for three small-molecules.

### 3.3. Interpreting the refined structures and crystal packing of the compounds

We have shown how the three datasets differ at the level of the refined molecular model (**Fig. 3a–c**) and its causal orientation within the unit cell (**Fig. 3d-i**). The refined structures are chemically reasonable and consistent with the expected molecular conformations; however, clear differences are observed in the ADPs of individual atoms. These models are represented by ellipsoids derived from the refined anisotropic displacement factors [6], and atoms with enhanced thermal motion; therefore, appear as expanded displacement ellipsoids.

**Figure 3.**
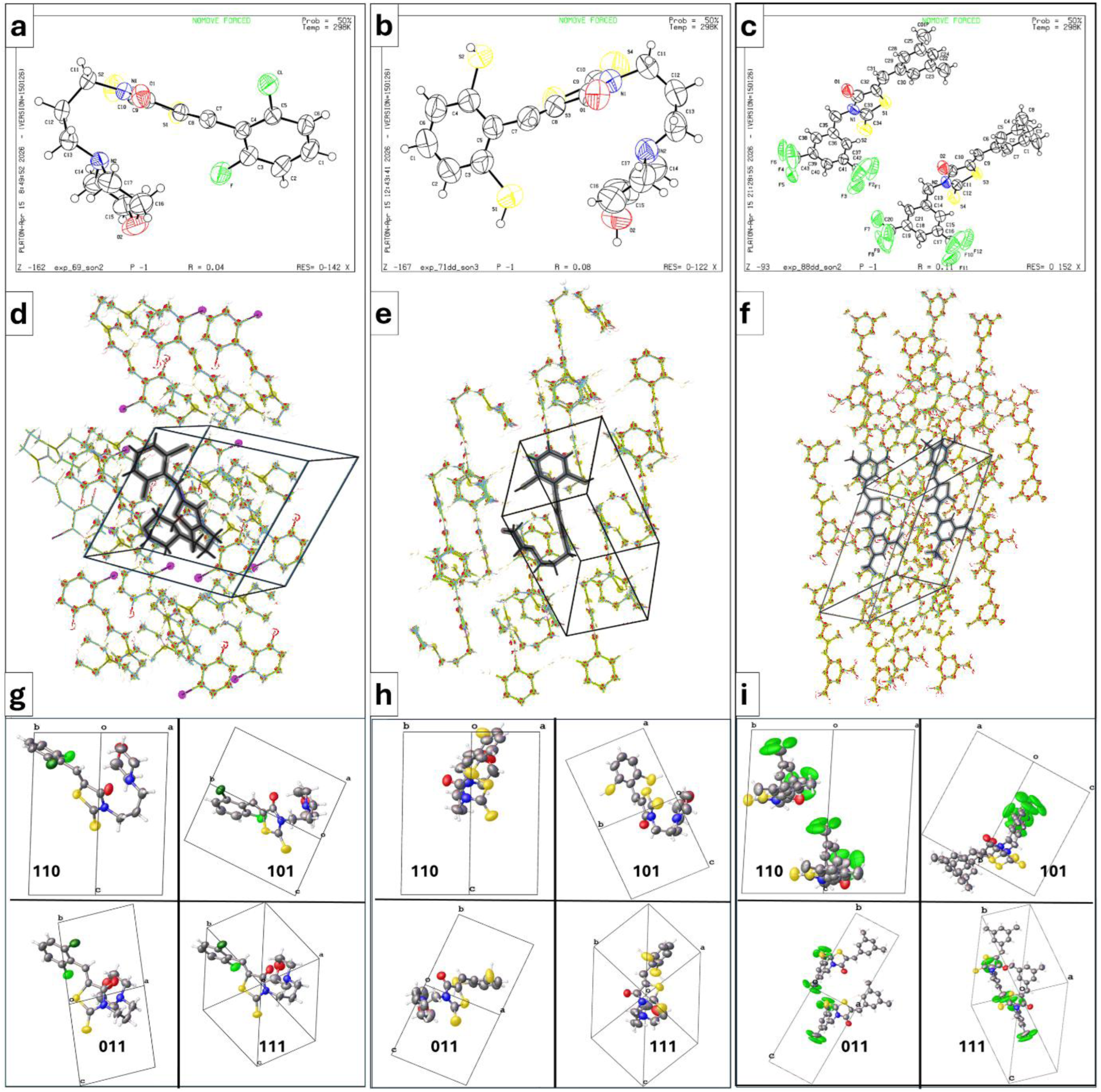
Analysis the refined molecular model of the compounds and their orientations within the unit cell. Comparative crystallographic zoom in c-5a, c-5p, and c-5b, showing (a-c) final refined molecular models with anisotropic displacement ellipsoids, (d-f) the global crystal-packing environments generated from the ASU and its symmetry mates, and (g-i) projections along the [110], [101], [011], and [111] zone axis to highlight differences in local packing density, molecular orientation, and structural complexity. In panel d-f, molecule in ASU is colored in black.

Accordingly, c-5a shows the well-ordered ellipsoid distribution overall, with the non-hydrogen atoms remaining comparatively compact and regular in shape (**Fig. 3a**). Likewise, c-5p remains structurally acceptable, but mean-square displacement amplitudes of atoms already display less uniform displacement ellipsoids than in c-5a, indicating a moderate increase in local motion and/or positional disorder (**Fig. 3b**). The effect is most evident in c-5b, where the fluorinated aryl group exhibits the apparent displacement ellipsoids (Ueq of F2 to F12 atoms is 0.212 Å² to 0.273 Å² in c-5b versus 0.081 Å² in single F of c-5a), making the F atoms the most prominent indicators of local disorder, enhanced positional displacement tensor, or unresolved structural ambiguity in the model (**Fig. 3c**). Such behavior of F is well-reported in small-molecule crystallography [11], where highly anisotropic displacement ellipsoids may reflect dynamic motion, static disorder, or limitations in the local structural model [6], [11]. The interpretation is also consistent with **Fig. 2**, in which c-5b gave the weakest agreement statistics and therefore behaved as the most challenging structure among the other three. This ambiguity may be further rationalized by the fact that the c-5b structure contains dissymmetric two identical molecules within ASU (Z′ = 2; Z′ is the symmetry-independent molecules in a unit cell), a feature that increases crystallographic complexity and can plausibly contribute to weaker refinement behavior [12].

Collective packing views highlight the local molecular environment in the crystal and show qualitative differences in the packing arrangement among the three datasets (**Fig. 3d–f**). Especially, the environment around c-5b appears denser, in line with its more complex structural and refinement issue (**Fig. 3f**). Additional viewing the crystal structures along the [110], [101], [011], and [111] zone axis provided detailed information on molecular geometry, orientation within the unit cell, and crystal packing (**Fig. 3g-i**) [13]. These views are not only alternatives of the same structural model; rather, they identified how the molecular conformation is oriented relative to the crystallographic axes and how the substituent(s) are distributed along different crystallographic directions [13]. Namely, the [110] zone axis reveals the overall molecular dimension within the cell, and for identifying whether substituent atoms are localized or radiated along a selected packing direction, while the [101] zone axis provides a complementary picture of the molecular tilt within the lattice and more clearly reveals substituents extending out of the main molecular plane. The [011] view helps assess if the packing is layered or offset relative to the unit-cell axes, whereas the [111] view provides a more oblique perspective and is often the clearest for visualizing the full three-dimensional distribution of the peripheral substituents.

Collectively, these comparisons suggest that the more highly substituted structures, especially c-5b, occupy the unit-cell volume more densely, with the substituted parts of the molecule extending further into the packing space (**Fig. 3f, 3i**). Therefore, those packing views indicate that higher substitution and complex ASU might be related to more difficult refinement.

### 3.4. Computed anisotropic displacement profiles for thermal motion analysis

To further compare the refined models at the atomic-displacement environment, ADP data were extracted from the CIF files and analyzed (**Table 3**). Accordingly, the anisotropic factors U11, U22, U33, U12, U13, and U23 were parsed, after which an approximate equivalent displacement parameter was calculated as Ueq = (U11 + U22 + U33)/3, and an anisotropy index was defined as the ratio of the largest to the smallest principal diagonal term, max(U11,U22,U33)/min(U11,U22,U33). The resulting plots therefore provide two complementary views of the refinement outcome: the Ueq/anisotropy profiles identify the most displacement-active atoms, whereas the heatmaps show how the individual 𝑈𝑖𝑗 factors are distributed across the structure.

**Table 3.**
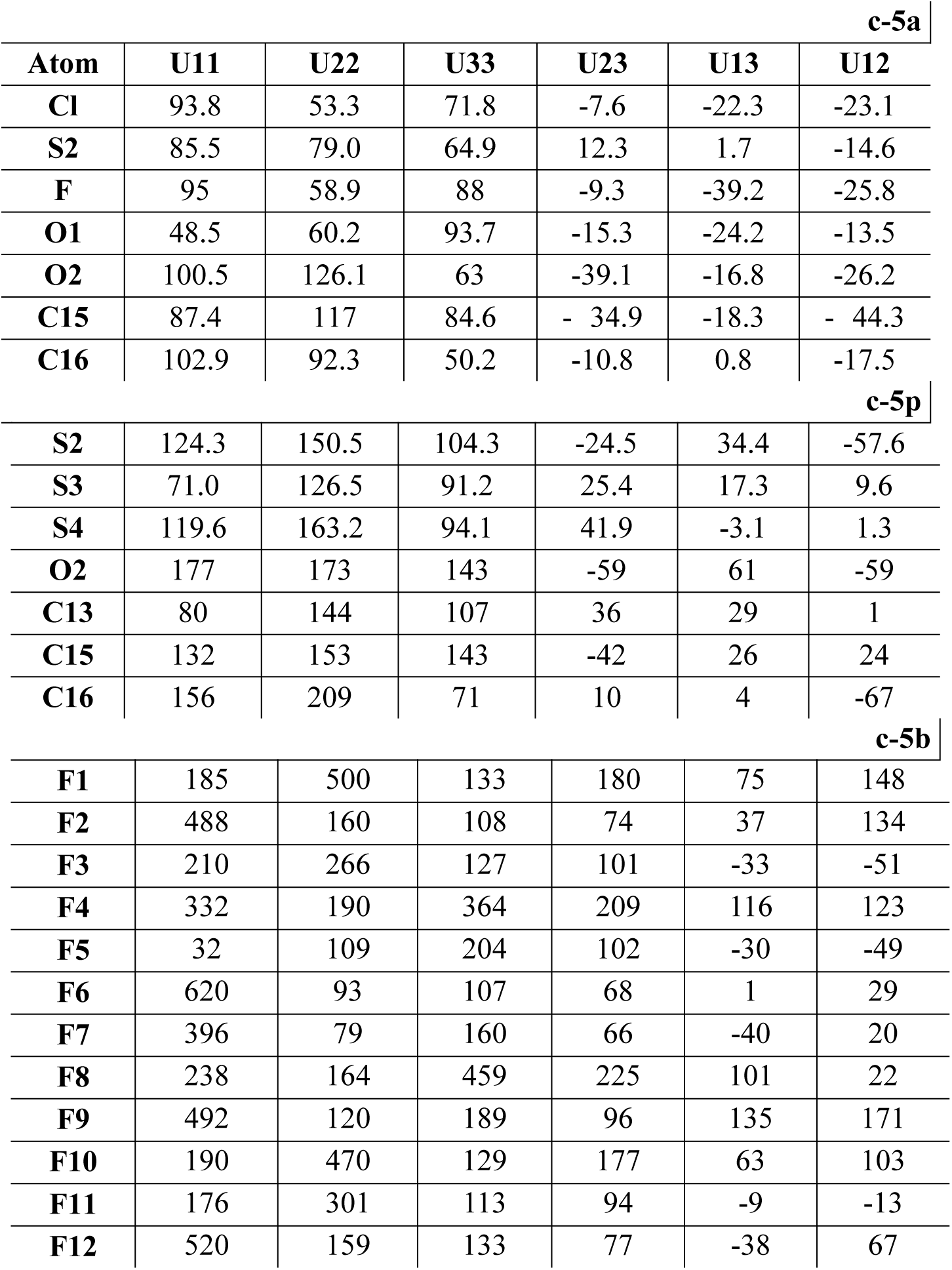
Anisotropic Displacement Parameters (Å^2^×10^3^) for three small molecules, respectively. The Anisotropic displacement factor exponent takes the form: [2π^2^[h^2^a*^2^U_11_+2hka*b*U_12_+…]5.

| **c-5a** |  |  |  |  |  |  |
| --- | --- | --- | --- | --- | --- | --- |
| **Atom** | **U11** | **U22** | **U33** | **U23** | **U13** | **U12** |
| **Cl** | 93.8 | 53.3 | 71.8 | -7.6 | -22.3 | -23.1 |
| **S2** | 85.5 | 79.0 | 64.9 | 12.3 | 1.7 | -14.6 |
| **F** | 95 | 58.9 | 88 | -9.3 | -39.2 | -25.8 |
| **O1** | 48.5 | 60.2 | 93.7 | -15.3 | -24.2 | -13.5 |
| **O2** | 100.5 | 126.1 | 63 | -39.1 | -16.8 | -26.2 |
| **C15** | 87.4 | 117 | 84.6 | - 34.9 | -18.3 | - 44.3 |
| **C16** | 102.9 | 92.3 | 50.2 | -10.8 | 0.8 | -17.5 |
| **c-5p** |  |  |  |  |  |  |
| **S2** | 124.3 | 150.5 | 104.3 | -24.5 | 34.4 | -57.6 |
| **S3** | 71.0 | 126.5 | 91.2 | 25.4 | 17.3 | 9.6 |
| **S4** | 119.6 | 163.2 | 94.1 | 41.9 | -3.1 | 1.3 |
| **O2** | 177 | 173 | 143 | -59 | 61 | -59 |
| **C13** | 80 | 144 | 107 | 36 | 29 | 1 |
| **C15** | 132 | 153 | 143 | -42 | 26 | 24 |
| **C16** | 156 | 209 | 71 | 10 | 4 | -67 |
| **c-5b** |  |  |  |  |  |  |
| **F1** | 185 | 500 | 133 | 180 | 75 | 148 |
| **F2** | 488 | 160 | 108 | 74 | 37 | 134 |
| **F3** | 210 | 266 | 127 | 101 | -33 | -51 |
| **F4** | 332 | 190 | 364 | 209 | 116 | 123 |
| **F5** | 32 | 109 | 204 | 102 | -30 | -49 |
| **F6** | 620 | 93 | 107 | 68 | 1 | 29 |
| **F7** | 396 | 79 | 160 | 66 | -40 | 20 |
| **F8** | 238 | 164 | 459 | 225 | 101 | 22 |
| **F9** | 492 | 120 | 189 | 96 | 135 | 171 |
| **F10** | 190 | 470 | 129 | 177 | 63 | 103 |
| **F11** | 176 | 301 | 113 | 94 | -9 | -13 |
| **F12** | 520 | 159 | 133 | 77 | -38 | 67 |

Accordingly, the anisotropy profile of c-5a remains relatively low and compact overall with only a few distinct maxima, notably around O1 (1.93), O2 (2.00), and C16 (2.05), indicating anisotropic displacement is present but remains relatively localized. The heatmap shows moderate variation with the largest intensity concentrated mainly in the diagonal terms (U11, U22, U33) and relatively modest off-diagonal components (U12, U13, U23), which are consistent with the well-ordered ellipsoid distribution of c-5a, reflecting the most internally coherent model among the three ones at the ADP level (**Fig. 4a**).

**Figure 4.**
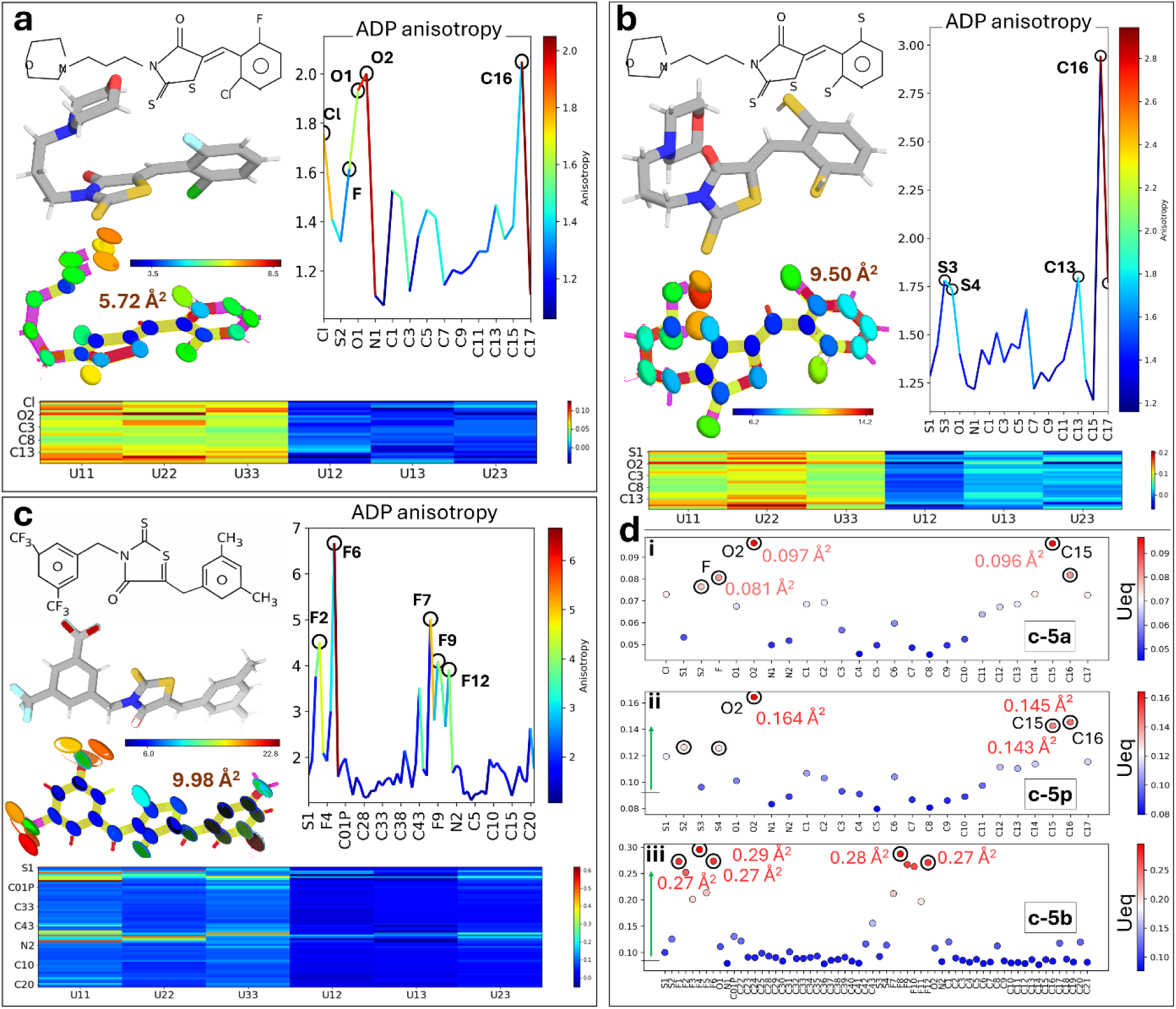
Comparative ADP analysis of c-5a, c-5p, and c-5b. Panels a-c present the molecular view, ellipsoid representation, atom-resolved anisotropy profile, and 𝑈𝑖𝑗 heatmap for each structure. (a) c-5a shows the most compact displacement pattern, (b) c-5p exhibits moderately increased local anisotropy, and (c) c-5b displays the strongest localized deviations, within the fluorinated aryl moiety. (d) Ueq scatter plots, confirming that the displacement pattern in c-5b is dominated by local displaced outlier atoms rather than by global elevation across the entire structure. The green arrow indicates that the y-axis maximum increases from c-5a to c-5p and c-5b, reflecting progressively larger Ueq ranges in the latter two structures.

The same analysis of c-5p indicates a moderately higher maximum displacement heterogeneity than c-5a, with prominent contributions from S3 (1.78), S4 (1.73), C13 (1.80), and especially C16 (2.94), suggesting that anisotropic displacement in c-5p is still localized, but is more pronounced than in c-5a, particularly at C16. The heatmap is still dominated by the diagonal 𝑈𝑖𝑖 factors but displays greater variation than c-5a. Thus, c-5p appears to retain an acceptable overall model quality with a larger number of local fluctuations (**Fig. 4b**).

The strongest deviations are observed for c-5b (**Fig. 4c**). Here, the anisotropy profile shows sharper and larger peaks than the other two, with the highest maxima related to the F2 (4.52), F6 (6.67), F7 (5.01), F9 (4.1), and F12 (3.19) atoms, indicating the fluorinated aryl moiety is the main source of possible structural ambiguity in c-5b. This is already directly supported by the tabulated values (**Table 3**); the Ueq for the F atoms in c-5b are apparently larger (0.212 Å² to 0.293 Å²) than the single F in c-5a (0.081 Å²), in line with ellipsoid representations. Even if the heatmap of c-5b is further consistent with localized enhancement of ADP factors, when combined with the Ueq scatter profile (**Fig. 4d**), the global displacement maxima in c-5b are not extremely elevated (Fig. 4c, 4diii; see the value for blue dots & gradually increase y-value marked by green arrow) relative to the other two, but are instead dominated by a limited number of high-displacement outliers (F alone).

In sum, the Ueq scatter of c-5a in **Fig 4di** shows modestly elevated Ueq values with the principal maxima localized at O2 (0.097 Å^2^), C15 (0.096 Å^2^), and F (0.081 Å^2^); whereas the displacement of c-5p becomes more pronounced, with O2 (0.164), C15 (0.145), and C16 (0.143∼) defining the main maxima (**Fig 4dii**). Collectively, c-5b shows well-supported different anisotropic distribution, characterized by higher outliers clustered in the fluorinated aryl moieties (**Fig 4diii**; ∼0.27-0.29 Å^2^), which is line with the structural and dynamic observations as mentioned above.

## 4. Discussion

The present study shows that the laboratory in-house diffractometer Turkish Light Source can produce publication-quality small-molecule structures when data collection, reduction, and refinement are performed carefully. This is particularly important for Türkiye, where modern in-house SCXRD source has only recently become available [4]. Early work from the same platform has already demonstrated its value for high-resolution structural analysis and for strengthening local crystallographic capacity [4], [5], [14]. The successful structure determination of compounds 5a, 5p, and 5b [2], [7], [8] indicates that this *boutique facility* is suited not only for proof-of-principle measurements but also for informative small-molecule studies. The workflow used here, crystal selection, data collection and reduction, generation of the *.ins file, refinement in Olex2 suite, and deposition of the final *.cif files in the Cambridge Crystallographic Data Centre (CCDC), therefore provides a coherent and reproducible pipeline for standard small-molecule structure determination (**Fig. 2**).

All compounds crystallized in the triclinic centrosymmetric space group P-1 and yielded reliable molecular models, although the quality of the datasets differed clearly (**Table 2**). c-5a gave the best ordered structure, c-5p remained reliable but somewhat less internally consistent, and c-5b demonstrated to be the most difficult to refine (**Fig. 2c**). This is supported by our independent observations: c-5a showed lower Rint values and stronger signal-to-noise, c-5p displayed intermediate behavior, and c-5b gave the weakest merging and refinement statistics together with higher SSE values and a more challenging refinement method (**Fig. 2d-f**). Because the same available experimental and computational pipeline was applied throughout, these differences might most reasonably be attributed to intrinsic structural characteristics of the crystals rather than variation in the experimental procedure.

The structural models help explain these differences; c- 5a and c-5b have the simplest substitution and show compact, regular displacement ellipsoids, consistent with relatively well-ordered structure (**Fig. 3a-b**). Conversely, c-5b is both more highly substituted and more complex crystallographically, since it contains two symmetry-independent molecules in the asymmetric unit (Z′ = 2) (**Fig. 3c,f,i**). This combination likely contributes to the problem of refinement (**Fig. 2ci**)

The anisotropic displacement analysis further supports our understanding. Displacement tensors of c-5a are compact and relatively isotropic, while c-5p becomes localized anisotropy, especially around selected S and C atoms (**Fig. 4a-b**). Interestingly, the largest anisotropy maxima of c-5b are focused on the fluorinated aryl groups, where the F atoms show higher Ueq values than the single F in c-5a (**Fig. 4c**). Suggests that the main crystallographic dilemma in c-5b arises from a limited group of atoms with localized ambiguity rather than from global disorder across the entire structure. Such an interpretation is agreeing with established crystallographic observations, where enlarged displacement ellipsoids often reflect enhanced thermal motion, unresolved static disorder, or local limitations of the structural model [6], [11]. The concentration of these effects in the fluorinated moiety is also chemically plausible, since heavily fluorinated peripheral groups often show elevated apparent ADPs [11] when closely related orientations or librational motions cannot be fully resolved.

Crystal packing of three provides further explanation: c-5b appears to occupy the unit-cell volume more densely than the other two, with substituents extending further into the intermolecular space and increasing steric congestion (**Fig. 3f**). This may promote local conformational heterogeneity within the lattice. The presence of two independent molecules in the asymmetric unit is likewise consistent with the prominent displacement heterogeneity (**Fig. 3c**) commonly observed in Z′ > 1 structures [12]. Inversely, the more compact packing of c-5a and c-5p agrees well with their more stable refinement process (**Fig. 2ci-ii**), suggesting that substituent complexity and packing density all contribute to the refinement tribulation of c-5b.

The results also highlight both the strengths and the limitations of ambient-temperature home-source crystallography. Data collected at 298 K with a laboratory Cu Kα source were acceptable to solve and refine all three compounds and to produce chemically reasonable structures for deposition in the CCDC [15]. This means an important useful crystal structure data for the newly established national facility and demonstrates the value of the Turkish home-source for standard small-molecule crystallography. Additionally, the elevated atomic displacement parameters observed in c-5b suggest that ambient-temperature structure may retain dynamic contributions to atomic motion that could be reduced under cryogenic conditions. A useful next step would therefore be to compare ambient- and low-temperature datasets for the same crystals, particularly for c-5b, to determine more clearly between thermal motion and static disorder in structurally ambiguous regions.

Overall, our results support three main conclusions; ***(i)***, the in-house TLS is robust for reliable small-molecule structure determination and complete pipeline from diffraction to CCDC deposition. ***(ii)***, among the three compounds studied, c-5a is the most ordered followed by c-5p, and c-5b is the most challenging due to localized displacement in its fluorinated aryl group and its Z′ = 2 asymmetric unit. ***(iii)***, our anisotropic displacement analysis is useful for determining the localized displacement effects from global disorders. Collectively, these findings show that careful home-source crystallography, combined with further displacement analysis, can both confirm structure and provide insight into the sources of refinement limitations in small-molecule SCXRD.

## Conclusion

This study reveals a modern in-house laboratory diffractometer, presented by the Turkish Light Source, can reliably produce publication-quality small-molecule crystal structures when combined with a consistent data-processing and refinement pipeline. The successful structure determination of three rhodanine-based domestic compounds (c-5a, c-5p, and c-5b) supports the efficacy of this end-to-end pipeline, from data collection to CCDC deposition. Although all datasets were collected and processed under identical conditions, the compounds showed notably diverse crystallographic surfaces. The observed movement (c-5a > c-5p > c-5b) is best described by disparities in intrinsic structural disorder rather than by crystallographic limitations. Indeed, the increased refinement ambiguity related to c-5b seems to be caused by its higher degree of substitution, densely packing space, and the presence of dissymmetric molecules (Z′ = 2). Notably, anisotropic displacement analysis provides a useful diagnostic perspective on refinement quality, suggesting that refinement ambiguity is more strongly related to localized maxima in the displacement within the fluorinated aryl moiety of c-5b rather than to global disorder of the entire structure. Combined with computational analysis, these findings indicate that in-house SCXRD can serve not only as a reliable strategy for small-molecule structure determination but also help explain which structural characteristics are liable to refinement tribulations. Incorporating low-temperature data collection in future studies may further clarify the comparative contributions of dynamic motion and static disorder, particularly in highly substituted systems.

## Data Availability

The crystallographic data for compounds 5a, 5p, and 5b have been deposited with the Cambridge Crystallographic Data Centre (CCDC) under deposition numbers 2546382, 2546533, and 2546833, respectively. These data can be obtained free of charge from the Cambridge Crystallographic Data Centre via www.ccdc.cam.ac.uk/data_request/cif

## Author Contributions

The analyte was synthesized by A.M. Small-molecule SCXRD setup was installed by E.A. and data collection, processing, and further analysis were performed by E.A. under the supervision of A.M. All authors have read and approved the final version of the manuscript.

## Acknowledgements

The authors gratefully acknowledge Şaban Tekin, Jakub Wojciechowski, and Ahmet Katı for their invaluable technical and administrative support in facilitating access to and use of the Turkish Light Source.

